# Latitude, not geography, globally structures *Oscheius tipulae* into three deeply divergent lineages

**DOI:** 10.64898/2026.06.26.734863

**Authors:** Dahyun Byeon, Daisy S. Lim, Junho Lee

## Abstract

Free-living nematodes are among the most abundant animals on Earth and play critical ecological roles in soil ecosystems. However, the global population structure and evolutionary history of most species remain poorly understood. Here, we analyzed genome-wide variation in *Oscheius tipulae* using whole-genome sequence data from 31 isolates, including 28 publicly available genomes and three newly collected strains from Korea. Population structure analyses, phylogenomic inference, and ancestry estimation consistently identified three deeply divergent lineages. These analyses did not detect admixture among lineages and collectively supported a predominantly tree-like evolutionary history. Notably, the lineages were structured by latitude rather than geographic proximity. Isolates from similar latitudinal zones clustered together regardless of continental origin, forming three major groups: northern mid-latitude (NML), low-latitude (LL), and southern mid-latitude (SML). This pattern indicates that the lineages have maintained largely independent evolutionary trajectories over extended timescales despite the potential for long-distance dispersal. Furthermore, environmentally associated variants showed significant differentiation among lineages, indicating that environmental selection may contribute to the maintenance of this latitudinally structured diversity. Our results reveal unexpectedly deep global divergence within *O. tipulae*, and highlight the importance of ecological divergence and long-term lineage retention in shaping the global diversity of this group.

## Introduction

Free-living nematodes are among the most abundant and widely distributed animals on Earth, inhabiting diverse terrestrial and aquatic environments including soils, decomposing organic matter, and microbial-rich substrates (Schratzberger et al., 2019; van den Hoogen et al., 2019). They can also function as pioneer organisms that rapidly colonize habitats during early stages following disturbance (Ptatscheck et al., 2015; Ptatscheck & Traunspurger, 2014). In soil ecosystems, nematodes play important ecological roles as decomposers and regulators of microbial communities, contributing to nutrient cycling and ecosystem functioning. Because of their high population densities and ecological ubiquity, nematodes are often considered key components of the soil food web and valuable model organisms for studying ecological and evolutionary processes (Ferris et al., 2001; Gebremikael et al., 2016; Neher, 2010).

Despite their ecological importance, the large-scale population structure and dispersal patterns of many free-living nematodes remain poorly understood. Most nematodes have limited intrinsic mobility due to their small body size, suggesting that their active dispersal capacity may be restricted, particularly over large spatial scales. Consistent with this expectation, low-dispersal small animals often exhibit isolation by distance (IBD), in which genetic differentiation increases with geographic distance as spatially limited dispersal restricts gene flow among populations (Wright, 1943). This pattern has been documented across a wide range of low-vagility taxa (Blouin et al., 1995; Cabe et al., 2007; Pfenninger et al., 1996). However, several studies have shown that nematodes can also be transported passively by external vectors such as wind, insects, or animals, raising the possibility of dispersal across much broader geographic ranges than expected from active movement alone (Ptatscheck & Traunspurger, 2020). Experimental and field studies have shown that living nematodes can be transported through the atmosphere and remain viable after aerial dispersal, suggesting that wind-mediated transport may represent one potential mechanism contributing to their geographic distribution (Ptatscheck et al., 2018).

The soil nematode *Oscheius tipulae* (Rhabditida: Rhabditidae) has emerged as an attractive system for studying population genetics and evolutionary processes in free-living nematodes (Baïlle et al., 2008; Félix, 2006). Similar to the model species *Caenorhabditis elegans*, *O. tipulae* reproduces primarily through self-fertilizing hermaphrodites with occasional males (Félix, 2006; Félix et al., 2000; Kiontke & Fitch, 2005). However, in contrast to *C. elegans*, *O. tipulae* exhibits a broader ecological distribution and has been isolated from diverse environments including soil, leaf litter, and decomposing plant material across multiple continents (Baïlle et al., 2008; Félix & Duveau, 2012; Sloat et al., 2022; Tintori et al., 2024). Previous studies have reported that *O. tipulae* maintains substantially higher levels of genetic diversity than other self-fertilizing nematodes such as *C. elegans* and *C. briggsae*, suggesting that *O. tipulae* may represent a useful system for investigating patterns of genetic diversity and population structure in free-living nematodes (Baïlle et al., 2008).

Although several studies have characterized genetic variation in *O. tipulae*, most have relied on limited genetic markers or geographically restricted sampling, leaving the global population structure of this species largely unresolved (Baïlle et al., 2008; Tintori et al., 2024). As a result, it remains unclear how genetic variation is organized across its worldwide distribution. Given the limited intrinsic mobility of nematodes but their potential for passive dispersal, *O. tipulae* provides an ideal system to investigate how dispersal processes and demographic history interact to shape genetic structure at a global scale.

In this study, we investigated the global population structure of *O. tipulae* using whole-genome sequencing data from 31 strains, including 28 publicly available isolates and three newly collected strains from South Korea. Through population genomic analyses of genome-wide variation, phylogenomic relationships, and ancestry patterns, we identified three deeply divergent lineages with no signs of admixture, supporting a predominantly tree-like evolutionary history. These lineages were more strongly associated with latitude rather than geographic proximity, revealing an unexpected global population structure that transcends major geographic barriers. Our results suggest that ecological factors and long-term lineage persistence, potentially shaped by environmental selection and large-scale dispersal processes, have played important roles in the global distribution and evolutionary history of this species.

## Results

### Global *O. tipulae* Populations Form Three Distinct Genetic Clusters

To investigate global population structure in *Oscheius tipulae*, we analyzed whole-genome sequencing data from 31 isolates, including 28 publicly available genomes (Dockendorff et al., 2022; Tintori et al., 2024) and three newly collected Korean samples (Supplementary Table S1).

Principal component analysis (PCA) based on genome-wide biallelic SNPs revealed three clearly separated clusters along the first two principal components. PC1 and PC2 explained 17.97% and 16.14% of the total genetic variance, respectively (Fig. 1a). Cluster1 and cluster2 were primarily separated along PC1, whereas cluster3 was clearly distinguished from the other two groups along PC2. Samples within each cluster appeared to be distributed along a gradual cline rather than forming tightly compact subgroups; however, despite this within-cluster continuity, the three clusters remained clearly separated with no apparent overlap among groups, supporting pronounced genetic structure. Given the predominantly selfing mating system of *O. tipulae*, which can result in extensive non-random associations among loci, we further evaluated the robustness of this pattern using SNP datasets pruned for linkage disequilibrium (LD). LD pruning was performed to remove highly correlated markers and retain a set of approximately independent SNPs, thereby minimizing the potential influence of redundant genomic signals and reducing bias in population structure inference. The same three major clusters were consistently recovered under both stringent (r² = 0.1) and permissive (r² = 0.8) LD-pruning thresholds (Supplementary Fig. S1). These results indicate that the observed population structure is robust to LD pruning and is unlikely to be driven solely by non-independence among linked markers.

**Fig. 1.**
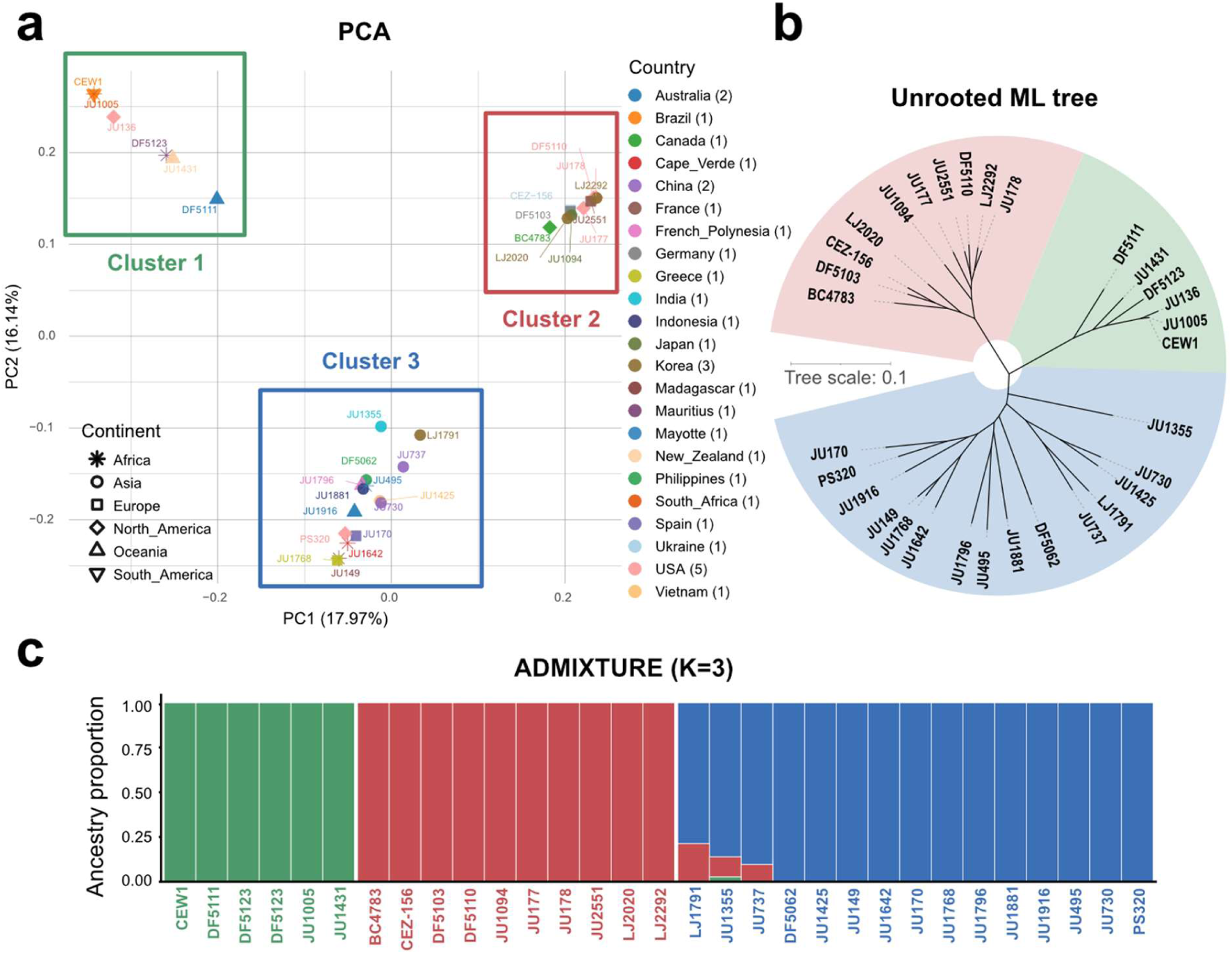
Global population structure of *O. tipulae*. **a,** Principal component analysis (PCA) of 31 *O. tipulae* isolates based on genome-wide biallelic SNPs. The first two principal components explain 17.97% (PC1) and 16.14% (PC2) of the total genetic variance. Three clearly separated genetic clusters are observed. Cluster1 and cluster2 are primarily separated along PC1, whereas cluster3 is separated from the other two clusters along PC2. Each point represents an individual isolate and is colored by sampling location. **b,** An unrooted maximum-likelihood phylogenetic tree constructed from genome-wide SNP data. The phylogeny is consistent with the PCA, separating the isolates into three major groups, with the groups distinguished by relatively long internal branches. **c,** Population structure inferred using ADMIXTURE with K = 3. Each vertical bar represents an individual isolate and colors indicate ancestry proportions. Colors used for sample labels indicate the clusters defined in the PCA and phylogenetic analyses. The results show minimal admixture among the three major clusters, supporting the presence of three genetically distinct populations. Alt text: Principal component analysis and phylogenetic tree showing 31 *O. tipulae* isolates clustered into three distinct genetic groups by genomic single nucleotide polymorphisms, with population structure bar chart demonstrating minimal admixture between clusters.

The unrooted maximum-likelihood (ML) phylogeny was consistent with the clustering pattern observed in the PCA, separating the isolates into three major groups (Fig. 1b). Isolates belonging to the same PCA cluster were also grouped closely together in the phylogeny, and the three groups were separated by relatively long internal branches. These results indicate that the isolates exhibit a clear genetic structure and are separated into three distinct groups. A rooted ML phylogeny was additionally reconstructed using *O. onirici* (Abolafia & Peña-Santiago, 2019; Torrini et al., 2015) as an outgroup to assess the evolutionary relationships among the three major groups (Supplementary Fig. S2). While the rooted tree consistently recovered the same three clusters identified by the PCA and unrooted phylogeny, it did not provide sufficient resolution to infer the branching order among these clusters.

Individual ancestry inference using ADMIXTURE (Alexander et al., 2009), a model-based approach for estimating genetic ancestry proportions, further supported K = 3 as the most likely number of genetic clusters based on cross-validation error (Supplementary Fig S4). Although a small number of samples showed minor proportions of ancestry from another cluster, most isolates were assigned almost entirely to a single ancestry component, indicating clear genetic differentiation among the three groups (Fig. 1c, Supplementary Fig. S3 and S4). Consistent with the PCA analysis, we further evaluated the effect of LD pruning by applying multiple pruning stringencies prior to ADMIXTURE inference. When a more permissive LD threshold was used (*r*² = 0.8), K = 3 was consistently recovered as the optimal number of clusters, whereas a more stringent threshold (*r*² = 0.1) resulted in K = 2 (Supplementary Fig. S3 and S4). However, under the stringent threshold, the cross-validation errors for K = 2 and K = 3 were nearly identical (1.106 and 1.117, respectively), suggesting that the preference for K = 2 was marginal rather than reflecting a substantially better fit to the data. Together with the consistent recovery of three clusters across PCA and phylogenomic analyses, these results support K = 3 as the most robust estimate of population structure.

Together with the PCA and phylogenetic analyses, the ADMIXTURE results consistently support that global *O. tipulae* populations are structured into three genetically distinct groups. The concordance among these independent analytical approaches indicates that the observed population structure is robust and reflects substantial genetic differentiation among the three major clusters.

### Population Clustering Is Strongly Associated with Latitude Rather Than Geographic Distance

When the genetic cluster assignments were mapped to the geographic origins of the isolates, the three genetic clusters exhibited a clear latitudinal distribution pattern (Fig. 2). Based on their geographic distribution, we designate cluster1, cluster2, and cluster3 as the Southern Mid-Latitude (SML), Northern Mid-Latitude (NML), and Low-Latitude (LL) groups, respectively. The NML group comprised isolates sampled from northern temperate regions of the Northern Hemisphere, the LL group included isolates collected primarily from tropical and low-latitude regions, and the SML group consisted of isolates from southern temperate regions in the Southern Hemisphere. Isolates sampled from similar latitudes but different continents clustered together genetically. For example, northern mid-latitude isolates from North America and Europe grouped within the NML cluster, whereas tropical isolates from Africa and Southeast Asia formed the LL cluster. Isolates from southern mid-latitude regions, including southern Africa and Oceania, belonged to the SML cluster. One exception to this pattern was an isolate collected from Hawaii, which clustered within the SML group despite being located in a low-latitude region. This pattern may reflect the unusually high genetic diversity reported from Hawaiian populations of free-living rhabditid nematodes, particularly *Caenorhabditis elegans* (Crombie et al., 2019; Cutter, 2006; Rockman & Kruglyak, 2009).

**Fig. 2.**
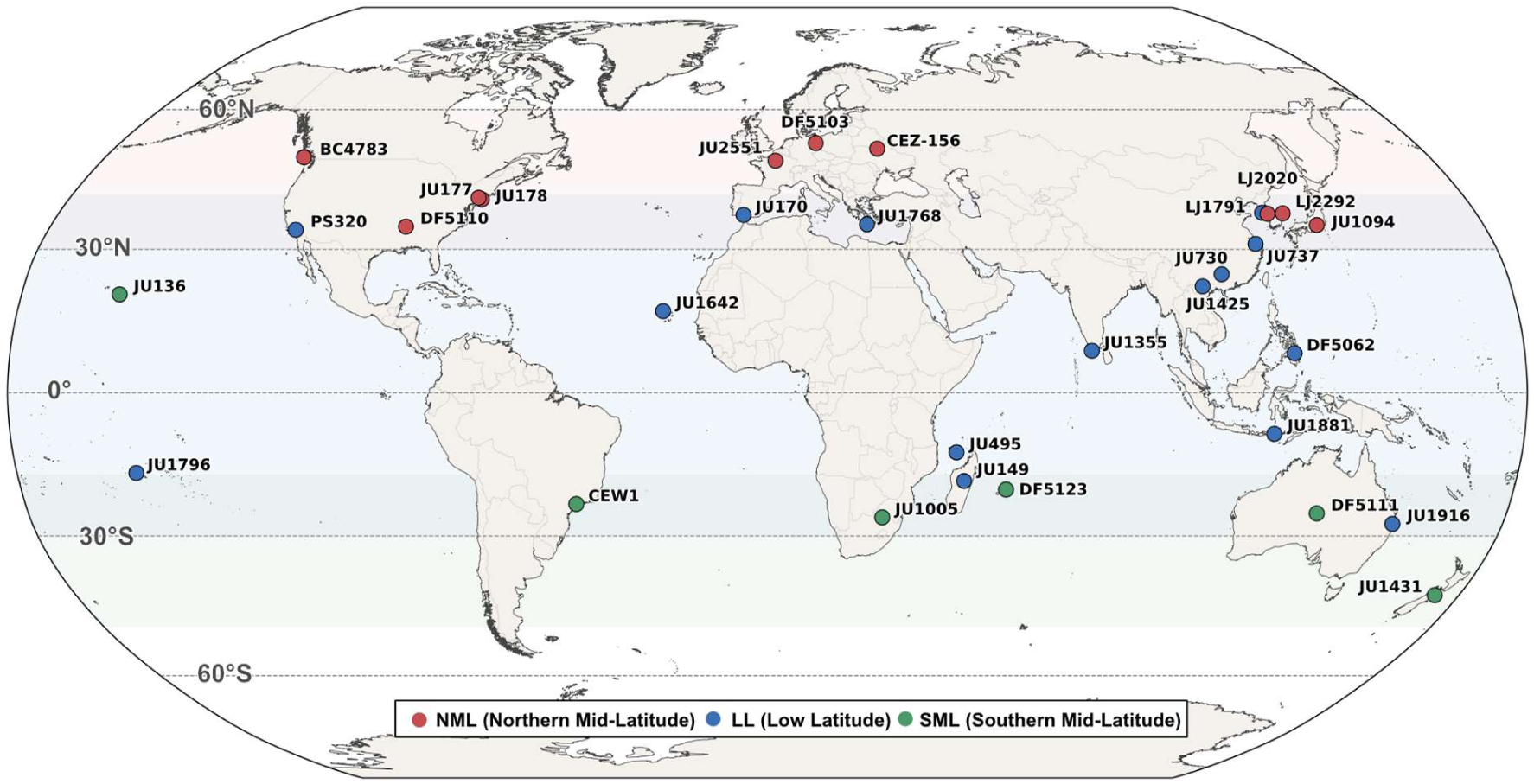
Latitudinal organization of global *O. tipulae*. Global sampling locations of the 31 *O. tipulae* isolates analyzed in this study. Isolates are colored according to the three genetic clusters identified by genome-wide analyses: Southern Mid-Latitude (SML, green), Low-Latitude (LL, blue), and Northern Mid-Latitude (NML, red). Shaded bands indicate the approximate latitudinal ranges occupied by each group. Despite occurring on different continents, isolates belonging to the same genetic cluster are largely restricted to similar latitudinal zones. Alt Text: World map displaying geographic distribution of 31 *O. tipulae* isolates color-coded by three latitudinal groups, with Northern Mid-Latitude in red, Low-Latitude in blue, and Southern Mid-Latitude in green.

To quantitatively evaluate the apparent latitudinal pattern observed in the geographic distribution of the genetic clusters, we performed Mantel tests (Mantel, 1967) comparing pairwise genetic distances with both geographic distances and latitudinal differences. The results showed that genetic distance was more strongly correlated with latitude than with overall geographic distance (Supplementary Fig. S5a, b). These results suggest that latitudinal factors, rather than geographic distance, better explain the global population structure of *O. tipulae*.

Notably, among the three newly collected Korean isolates, one clustered within the NML group, whereas the remaining two were assigned to the LL group. Despite their close geographic proximity, these isolates were assigned to different genetic clusters. This pattern may indicate that the Korean region lies near a potential boundary between the two genetic groups.

### Historical Relationships Among Clusters Show No Evidence of Gene Flow

To understand the historical processes underlying the observed latitudinal genetic structure, we investigated the evolutionary relationships among the three major genetic clusters and tested for evidence of post-divergence gene flow. First, admixture three-population (f3) statistics (Patterson et al., 2012; Reich et al., 2009) were used as a formal test for whether any cluster could be modeled as a mixture of the other two clusters. Significantly negative f3 values are expected when a target population has experienced admixture from the two reference populations. Second, outgroup f3 statistics (Patterson et al., 2012; Reich et al., 2009) were calculated to quantify the amount of shared genetic drift between pairs of clusters relative to an outgroup, thereby providing a measure of pairwise genetic affinity. Third, f4 statistics (Patterson et al., 2012; Reich et al., 2009) were used to distinguish alternative tree-like relationships among the three clusters and to test whether allele-frequency correlations were consistent with a simple bifurcating history or instead suggested excess allele sharing. Finally, TreeMix (Pickrell & Pritchard, 2012) was used to model genome-wide allele-frequency covariance in a graph-based framework and to evaluate whether adding migration edges improved the fit over a migration-free tree.

Admixture f3 statistics showed no evidence of admixture among the three major genetic clusters. Across all possible combinations of the three clusters as target and source populations, none yielded significantly negative f3 values (all f3 > 0; Z-scores ranging from 7.12 to 9.34; Table 1). These results indicate that the major genetic groups have remained largely genetically distinct following their divergence.

**Table 1.** Results of three-population (f3 statistics) tests for the three major *O. tipulae* clusters. Negative f3 values with |Z| ≥ 3 indicate evidence of admixture. None of the tested combinations produced significantly negative f3 values (all f3 > 0; Z > 7).

| Source 1 | Source 2 | Target | $f_3$ statistics | Standard error | Z-score | SNPs |
| --- | --- | --- | --- | --- | --- | --- |
| cluster1 | cluster2 | cluster3 | 0.195101 | 0.021244 | 9.184 | 1,629,619 |
| cluster2 | cluster3 | cluster1 | 1.261702 | 0.177175 | 7.121 | 1,629,619 |
| cluster3 | cluster1 | cluster2 | 0.76696 | 0.082118 | 9.34 | 1,629,619 |

Outgroup f3 statistics, calculated using *O. onirici* as the outgroup, revealed clear differences in shared genetic drift among the three latitudinal lineages. As expected, comparisons within each lineage showed higher levels of shared drift than comparisons between lineages. Among the between-lineage comparisons, NML and LL exhibited relatively higher shared drift than lineage pairs involving SML (Fig. 3a), indicating a closer genetic affinity between NML and LL.

**Fig. 3.**
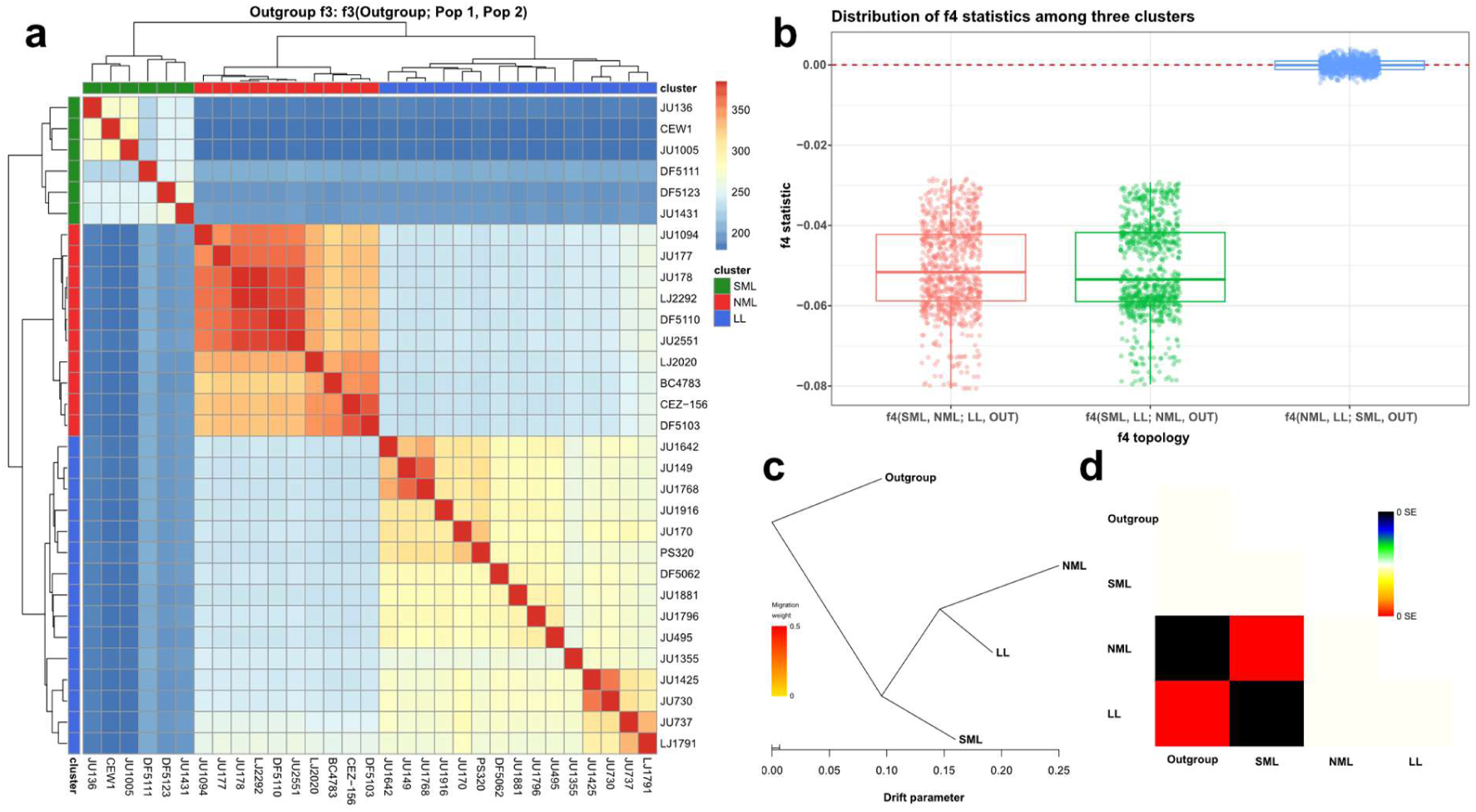
Historical relationships among latitudinal lineages support a predominantly tree-like evolutionary history. **a,** Heatmap of outgroup f3 statistics calculated using *Oscheius onirici* as the outgroup. Warmer colors indicate greater shared genetic drift. Comparisons within lineages show the highest shared drift, whereas among-lineage comparisons reveal stronger affinity between NML and LL than between either lineage and SML. **b,** Distribution of f4 statistics for alternative lineage relationships. Values for f4(NML, LL; SML, *O. onirici*) are centered near zero, whereas alternative topologies show systematic deviations from zero, supporting a tree-like relationship in which NML and LL form sister lineages relative to SML. **c,** TreeMix population graph inferred without migration edges (m = 0). The topology recovered NML and LL as sister lineages, with SML forming the earliest-diverging lineage among the three major groups. Branch lengths are proportional to genetic drift. **d,** Residual covariance matrix from the TreeMix model. Residuals are generally small, indicating that the migration-free tree adequately explains the major covariance structure among lineages, providing little evidence for substantial recent gene flow among lineages. Alt Text: Outgroup f3, f4 statistics and TreeMix analyses support a sister relationship between NML and LL, with SML as the earliest-diverging lineage. A migration-free tree adequately explains the major covariance structure among lineages.

We next calculated f4 statistics for the three lineages using all possible strain combinations. The distribution of f4(NML, LL; SML, *O. onirici*) was centered near zero, whereas the alternative configurations showed systematic deviations from zero (Fig. 3b). This pattern supports a largely bifurcating, tree-like relationship in which NML and LL are more closely related to each other than either is to SML, rather than indicating extensive post-divergence gene flow among lineages.

TreeMix, under a migration-free model (m = 0), supported the same tree-like relationship among the three major lineages inferred from the f-statistic analyses (Fig. 3c). Examination of the residual covariance matrix revealed no pronounced deviations from the fitted graph (Fig. 3d). Models incorporating one or two migration edges produced negligible improvements in log-likelihood compared with the migration-free model, and the inferred topology remained unchanged (Table 2). These results suggest that the major covariance structure among lineages is adequately explained by a tree-like model alone, without requiring admixture among the major lineages.

**Table 2.**
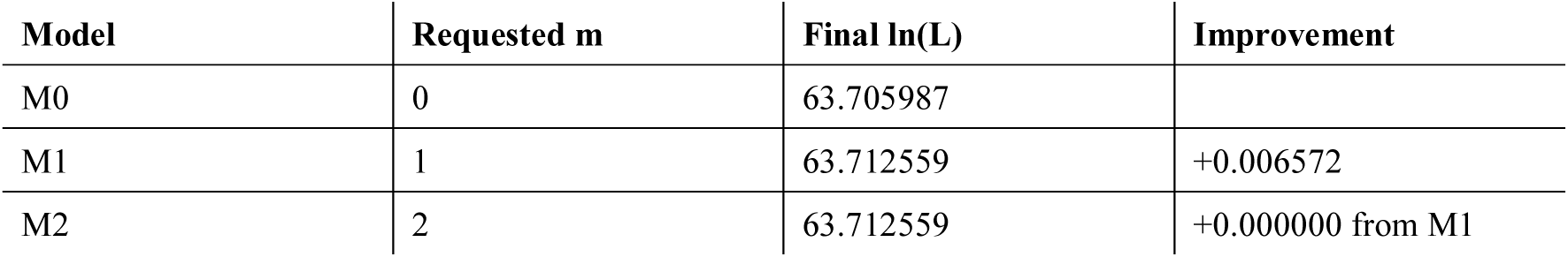
Comparison of TreeMix models with different numbers of migration edges. Final log-likelihood values for TreeMix models allowing 0, 1, or 2 migration edges. Incorporating a single migration edge resulted in only a marginal increase in log-likelihood (ΔlnL = 0.0066), whereas allowing a second migration edge provided no additional improvement.

Taken together, the concordant results obtained from multiple complementary approaches indicate that the genetic relationships among the major clusters are best explained by historical divergence rather than pervasive gene exchange. The consistency of these inferences across analytical methods highlights the stability of the underlying population structure and provides a coherent evolutionary context for the global distribution of genetic diversity in *O. tipulae*.

### Latitudinal groups exhibit distinct patterns of diversity and divergence

We next examined genome-wide patterns of genetic diversity and divergence among the three latitudinal lineages using nucleotide diversity (π), Hudson’s F_ST_, and absolute sequence divergence (D_XY_) (Hudson et al., 1992; Nei & Li, 1979). Nucleotide diversity was used to quantify standing genetic variation within each lineage, whereas F_ST_ and DXY were used to characterize genetic differentiation between lineages. F_ST_ is a relative measure that can be influenced by levels of within-lineage diversity, while D_XY_ reflects the absolute accumulation of sequence differences between lineages. Together, these statistics allow genetic differences among lineages to be evaluated from both within- and between-lineage perspectives, providing a more comprehensive view of population divergence.

Nucleotide diversity differed markedly among lineages. LL showed approximately two-fold higher diversity than either mid-latitude lineage (Table 3), indicating substantially greater standing genetic variation at low latitudes.

**Table 3.** Genome-wide estimates of nucleotide diversity (π), genetic differentiation (Hudson’s F_ST_), and absolute sequence divergence (D_XY_) among the three latitudinal lineages. Values on the diagonal represent nucleotide diversity (π) within each lineage. Values in the lower triangle represent pairwise Hudson’s F_ST_, and values in the upper triangle represent pairwise absolute sequence divergence (D_XY_). LL exhibited substantially higher nucleotide diversity than either mid-latitude lineage. Both pairwise Hudson’s F_ST_ and D_XY_ were lowest between NML and LL.

|  | NML | LL | SML |
| --- | --- | --- | --- |
| NML | 0.00495 | 0.01114 | 0.01149 |
| LL | 0.390 | 0.00859 | 0.01156 |
| SML | 0.599 | 0.449 | 0.00417 |

Genome-wide Hudson’s F_ST_ revealed pronounced genetic differentiation among lineages, with the highest value observed between the two mid-latitude lineages, followed by SML–LL and NML–LL (Table 3.).

In contrast, pairwise absolute sequence divergence (D_XY_) showed only modest variation among lineage pairs. The lowest D_XY_ value was observed between NML and LL, whereas slightly higher values were observed for NML–SML and SML–LL (Table 3.).

The partial discordance between F_ST_ and D_XY_ likely reflects differences in within-lineage diversity, particularly the elevated nucleotide diversity observed in LL. Nevertheless, both statistics support the same overall relationship among lineages, identifying NML and LL as the most closely related pair and SML as the most differentiated lineage- consistent with the tree-like history inferred from the f-statistic and TreeMix analyses.

### Population-specific differentiation and environmental associations identify candidate regions for local adaptation

Having characterized genome-wide patterns of differentiation shaped by demographic history and largely neutral evolutionary processes, we next investigated whether specific genomic regions showed signatures of population-specific divergence potentially associated with local adaptation and environmental variation.

To address this question, we quantified Population Branch Excess (PBE) (Yassin et al., 2016) across the genome for each latitudinal group. PBE measures allele-frequency differentiation along a focal population branch relative to the others, allowing the identification of genomic regions that show stronger population-specific divergence than expected from the genome-wide background. Such regions are potential candidates for lineage-specific selection or local adaptation. Using this approach, we identified genomic regions exhibiting elevated levels of population-specific allele-frequency differentiation in each latitudinal group.

We next evaluated whether these signals were associated with environmental variation. Based on the geographic coordinates of each sampling location, we extracted 20 environmental variables (e.g., temperature-and precipitation-related variables) and performed latent factor mixed model (LFMM) (Caye et al., 2019; Farmery et al., 2018) analyses to identify SNPs significantly associated with each environmental variable. We then intersected these LFMM candidate SNPs with the top 1% PBE regions identified for each population to detect genomic regions exhibiting both elevated population-specific differentiation and significant environmental associations. Genes located within these overlapping regions were then examined to identify candidate genes potentially underlying local adaptation to environmental variation. Candidate genes were annotated based on orthology to *C. elegans*. To distinguish putative *O. tipulae* genes from their *C. elegans* orthologs, candidate genes are referred to using the prefix “Oti-” followed by the corresponding *C. elegans* gene name unless otherwise noted. We highlight a subset of candidate genes for which functional information is available to facilitate interpretation of their associations with environmental variables and provide the full list in Supplementary Table S2-4.

The number and intensity of high-PBE peaks differed among the three population branches. High-PBE peaks were defined as genomic windows within the upper 1% of the empirical PBE distribution. The mean and median PBE values of these peaks were highest in NML (mean = 1.998, median = 1.896), followed by SML (mean = 1.767, median = 1.601), whereas LL exhibited substantially lower values (mean = 0.834, median = 0.713). These results indicate that population-specific differentiation is strongest in the two mid-latitude lineages and comparatively weaker in the low-latitude lineage (Fig. 4, Supplementary Table S2-4).

**Fig. 4.**
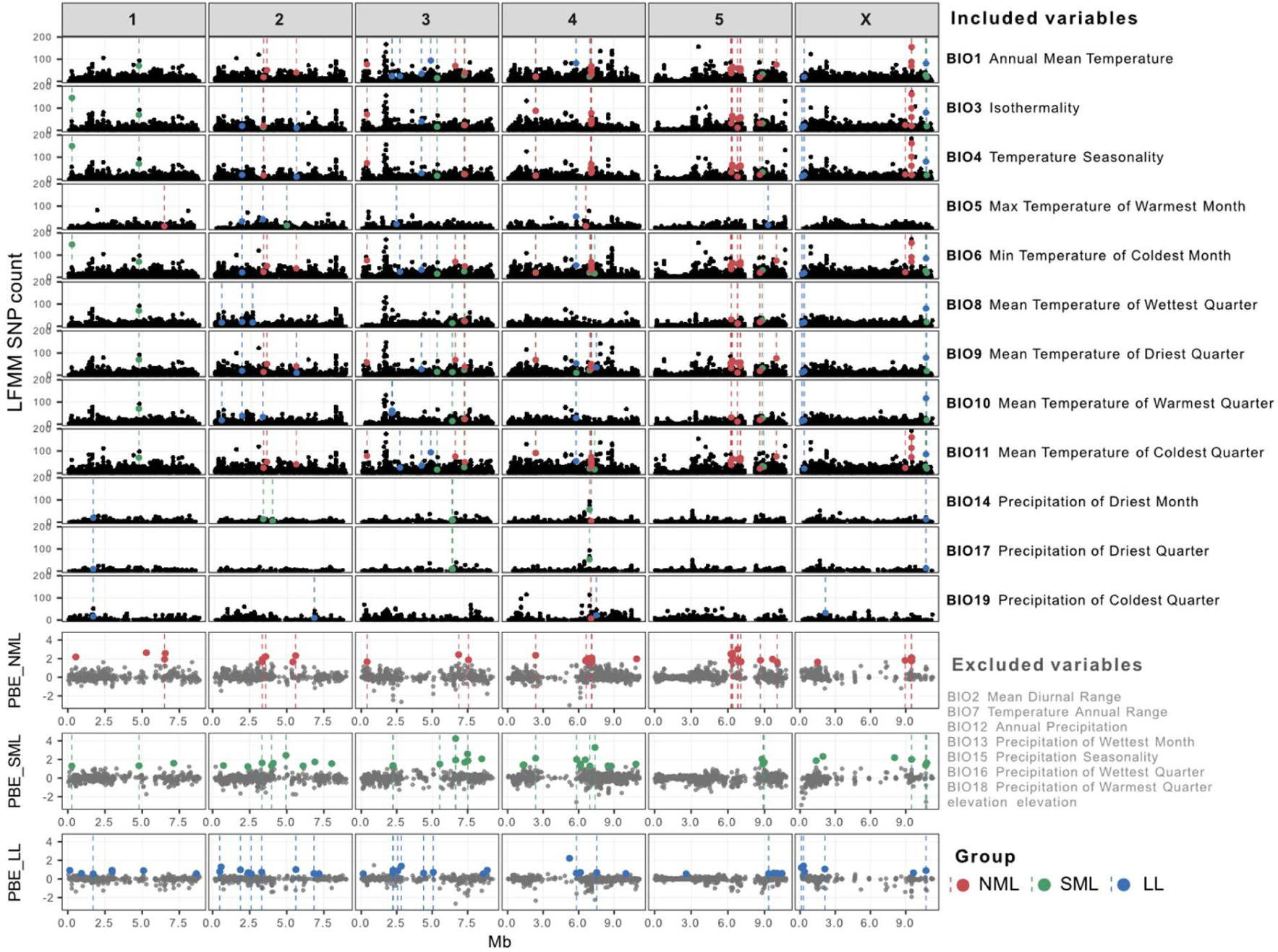
Integration of population-specific differentiation and environmental associations across the genome in *O. tipulae*. Genome-wide distribution of population-specific differentiation (PBE) and environmental associations (LFMM) across chromosomes. Each column represents a chromosome (1–5 and X), and genomic position is shown in megabases (Mb). Top panels, LFMM results showing the number of SNPs significantly associated with each environmental variable across genomic windows. Environmental variables are listed on the right. Bottom panels, PBE values for each population (NML, SML, LL), indicating population-specific excess allele frequency differentiation. Colored points represent windows exceeding the top 1% of the empirical PBE distribution. Vertical dashed lines indicate genomic regions where high PBE signals overlap with LFMM candidate regions. Included environmental variables are shown in black, while excluded variables (e.g., highly correlated variables or elevation) are shown in gray. Together, these results highlight genomic regions exhibiting both elevated population-specific differentiation and significant environmental associations, consistent with candidate loci underlying local adaptation. Alt Text: Genome-wide distribution of environmental associations and population-specific differentiation across five chromosomes and X.

The environmental associations of differentiated regions also differed among lineages (Fig. 5). Most overlap regions were associated with temperature-related variables, whereas precipitation-related associations were rarer. Temperature-associated overlaps were particularly abundant in NML, accounting for 55.1% of all temperature-related PBE–LFMM overlaps, compared with 23.9% in SML and 21.1% in LL. In contrast, precipitation-associated overlaps were distributed primarily between SML and LL, with only a small contribution from NML. These patterns suggest that distinct environmental factors are associated with lineage-specific differentiated regions across latitudinal groups.

**Fig. 5.**
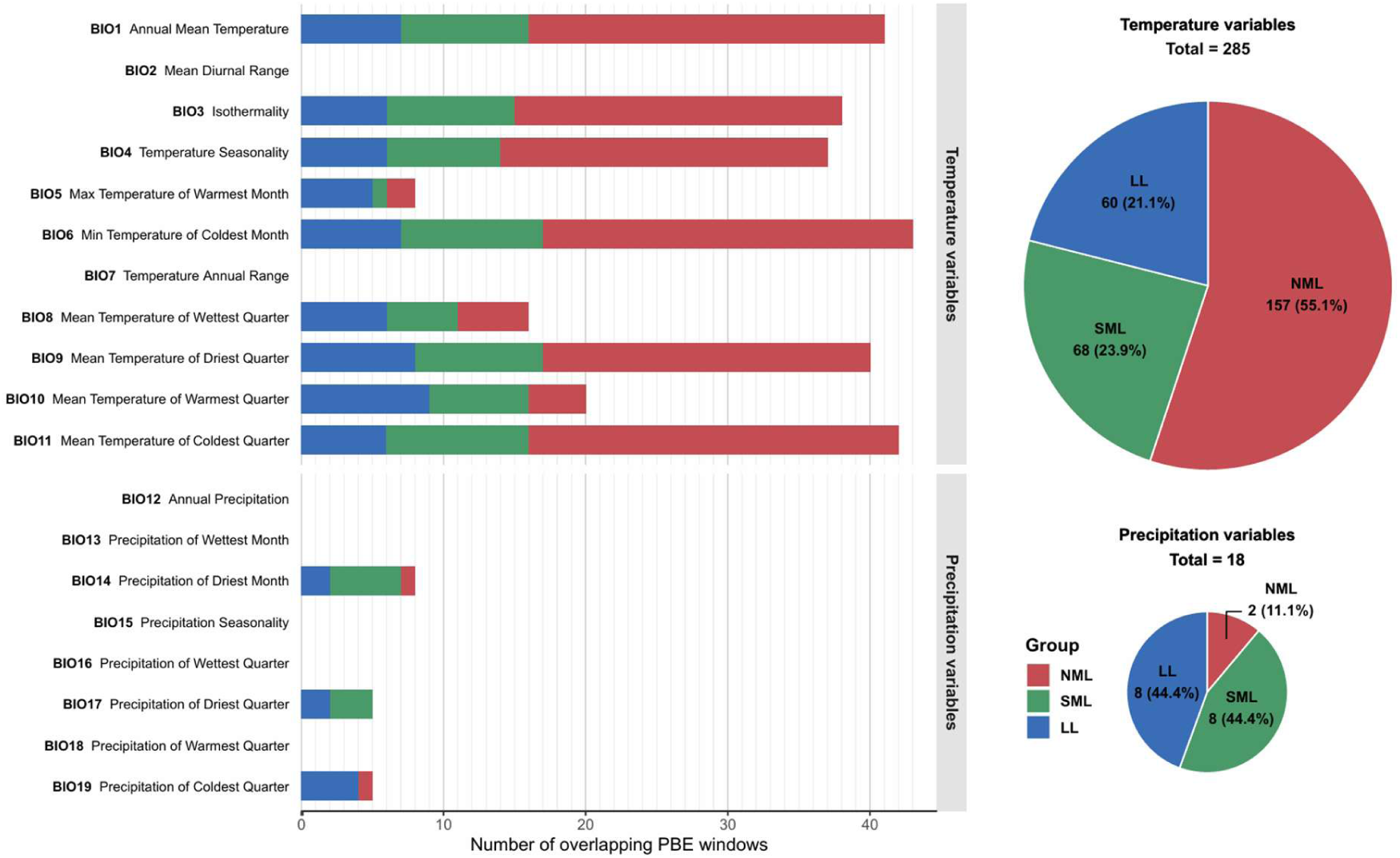
Distribution of PBE–LFMM overlaps across environmental variables and populations in *O. tipulae*. Number of genomic windows showing overlap between high PBE (top 1%) and significant LFMM signals across environmental variables. Bar plots show the number of overlapping windows for each environmental variable, partitioned by population (NML, SML, LL). Variables are grouped into temperature- and precipitation-related categories. Pie charts summarize the total number and proportion of overlaps contributed by each population for temperature (top) and precipitation (bottom) variables. Alt Text: Bar chart and pie charts showing overlap between population-specific differentiation and environmental associations.

The annotated candidate genes within these overlap regions also exhibited lineage-specific functional tendencies. In NML, several candidate genes were associated with sensory and neuronal signaling as well as proteostasis-related processes. Candidate genes identified in SML included genes associated with DNA repair, genome maintenance, redox-related processes, membrane transport, and RNA processing. In LL, candidate genes included genes associated with lipid metabolism, membrane or ion transport, and signaling pathways. Representative examples included *Oti-hlh-30* and *Oti-gcy-29* in NML, *Oti-xpd-1* and *Oti-trx-4* in SML, and *Oti-fat-2* and *Oti-best-9* in LL (Supplementary Tables S2–S4).

Together, these results indicate that a subset of population-specific differentiated regions also exhibit significant associations with environmental variables. While genome-wide analyses presented in the previous sections revealed strong genome-wide neutral divergence among the three latitudinal lineages, the combined PBE and LFMM analyses further demonstrate that differentiated genomic regions with putative functional roles are associated with distinct environmental variables and functional gene categories in each lineage.

## Discussion

In this study, we analyzed the global population structure of *Oscheius tipulae* using whole-genome sequencing data and genome-wide SNP analyses. Across multiple population genomic approaches, including principal component analysis, phylogenetic reconstruction, ancestry inference, and measures of genetic differentiation, we consistently identified three genetically distinct groups corresponding to Northern mid-latitude (NML), Low-latitude (LL), and Southern mid-latitude (SML) regions.

A prominent feature of this structure is its closer alignment with latitude than with geographic distance. Our results show that isolates from geographically distant regions but similar latitudes cluster together genetically, whereas geographically proximate isolates belonging to different latitudinal zones are assigned to distinct genetic groups.

The historical relationships among the three latitudinal groups were most consistent with a scenario of long-term divergence without detectable post-divergence admixture. Multiple complementary approaches converged on the same pattern: f-statistic analyses provided no evidence of substantial admixture among groups, while TreeMix indicated that genome-wide covariance could be explained without invoking migration events. Across these analyses, NML and LL consistently exhibited greater genetic affinity to one another than either did to SML, suggesting a shared history following separation from the southern mid-latitude lineage.

Patterns of genetic diversity and differentiation further refined this picture. The low-latitude lineage harbored substantially higher standing genetic variation than either mid-latitude group, whereas measures of population differentiation indicated pronounced genetic subdivision among all three lineages. Although relative differentiation (F_ST_) and absolute divergence (D_XY_) exhibited partially different patterns, both supported the same broad relationships recovered from the historical analyses. Together, these results suggest that contemporary population structure largely reflects the accumulation of divergence over extended evolutionary timescales rather than extensive secondary contact among lineages.

Genome-wide measures of differentiation revealed substantial divergence among all three latitudinal lineages, indicating long-term population subdivision across the global range of *O. tipulae*. F-statistic analyses and TreeMix consistently supported a tree-like relationship in which NML and LL are more closely related to each other than either is to SML. This topology was corroborated by both pairwise D_XY_ and genome-wide Hudson’s F_ST_ estimates, which identified NML and LL as the least divergent and least differentiated lineage pair.

In parallel, environmental association analyses identified signals consistent with population-specific responses at a subset of loci. By integrating Population Branch Excess (PBE) with LFMM analyses, we detected genomic regions that exhibit both elevated population-specific differentiation and significant associations with environmental variables. These candidate regions differed in their functional composition across populations, suggesting that distinct biological pathways may contribute to environmental responses in each group.

These findings build directly upon earlier population genetic studies of *O. tipulae*. Using AFLP markers, Baïlle et al. (2008) first demonstrated that global populations harbor substantial genetic diversity and are broadly structured into two major genetic groups associated with latitude. Our results not only corroborate those initial observations but substantially refine them: the increased resolution afforded by whole-genome sequencing and genome-wide SNP data revealed an additional level of population subdivision, resolving three well-supported latitudinal lineages (NML, LL, and SML) where previous marker-based analyses had identified only two major groups. Beyond refining population boundaries, the genome-wide dataset enabled the integration of complementary analytical approaches - including clustering analyses, phylogenetic reconstruction, ancestry inference, tests of population relationships and historical gene flow, and environmental association analyses - that were not feasible with earlier markers, together providing a far more detailed picture of how genetic diversity is partitioned across the global range of *O. tipulae*.

The magnitude and consistency of this structure across multiple genome-wide analyses indicate that NML, LL, and SML represent deeply differentiated evolutionary units rather than shallow population subdivisions, a degree of divergence that might, at first glance, raise the possibility of cryptic speciation. This interpretation is, however, explicitly contradicted by prior crossing experiments by Baïlle et al. (2008), in which reciprocal crosses among strains representing all three lineages produced viable and fertile offspring, confirming full reproductive compatibility under the biological species concept. These three lineages therefore constitute deeply diverged but reproductively cohesive units within a single biological species. A key question that follows is how such pronounced latitudinal structure has been maintained in a species capable of passive long-distance dispersal - a dispersal mode that would, in principle, be expected to continuously homogenize allele frequencies across geographic space.

One possible explanation is that long-distance dispersal, although evidently capable of connecting distant localities, may not occur randomly with respect to latitude. Passive dispersal vectors are often subject to directional biases imposed by prevailing physical gradients. For example, major components of the global wind belt system, such as the trade winds and westerlies, predominantly facilitate movement along east–west trajectories within similar latitudinal zones rather than across latitudes. If such biases generally promote dispersal within latitudinal bands rather than between them, the result would be asymmetric gene flow that maintains within-lineage cohesion while permitting prolonged divergence between lineages. Although LFMM and PBE analyses identified lineage-specific environmental associations at a subset of loci, the fact that all three lineages survive and reproduce normally under identical laboratory conditions suggests that the observed population structure does not arise from absolute physiological or ecological incompatibility among lineages. The maintenance of this structure is therefore more parsimoniously explained by structural constraints on dispersal opportunity itself than by hard ecological barriers imposed by environmental differences.

Notwithstanding these insights, several limitations warrant consideration. First, the number of isolates included in this study is relatively limited, and additional sampling across underrepresented regions will be necessary to further refine the global population structure of *O. tipulae*. Second, although the combination of PBE and LFMM analyses provides a useful framework for identifying candidate genomic regions, additional functional validation will be required to determine the precise roles of individual genes and variants.

## Materials and Methods

### Study samples and data sources

In this study, we analyzed whole-genome sequencing (WGS) data from a total of 31 *Oscheius tipulae* strains. Among these, 23 strains were obtained from a previously published dataset (Dockendorff et al., 2022), and 5 strains were obtained from another published study (Tintori et al., 2024) (Supplementary Table S1). The remaining three strains (LJ1791, LJ2020, and LJ2292) were newly collected from multiple locations in South Korea (Yeonpyeong Island, Gwanak Mountain, and Ulleung Island) and sequenced in this study. Metadata for all strains, including geographic origin, habitat, and sequencing information, are provided in Supplementary Table S1.

### Sample Collection and Whole-Genome Sequencing

Plant and soil samples were collected and put into plastic zipper bags or 50 mL Falcon tubes. Distilled water was added to each sample and mixed thoroughly, and aliquots of the resulting suspension were transferred dropwise onto standard *Caenorhabditis elegans* culture plates consisting of nematode growth medium (NGM) seeded with *Escherichia coli* OP50. Nematodes that migrated from the sample onto the bacterial lawn were collected the following day and transferred to fresh NGM plates. Androdioecious nematode strains capable of self-propagation from a single L1–L3 stage individual were maintained using the agar chunking method, which preserves both the nematode and its associated bacterial community.

Species identification was performed by PCR amplification of the 18S rDNA locus using primers nem1 and nem2 (Foucher & Wilson, 2002), followed by sequence similarity searches against the NCBI nucleotide database using BLAST (Altschul et al., 1990). For genomic DNA extraction, nematodes were harvested from culture plates as follows. LJ1791 worms were collected by thoroughly washing culture plates with distilled water, followed by eight consecutive washing steps consisting of centrifugation at 800 × g for 2 min, removal of the supernatant, and resuspension in fresh distilled water to minimize bacterial contamination. For LJ2020 and LJ2292, frozen stocks were thawed and propagated on NGM plates prior to harvesting. Worms were collected by adding distilled water to the plates and recovering the supernatant containing floating, actively swimming individuals. Genomic DNA was extracted using the Qiagen Puregene Cell and Tissue Kit (Qiagen, Cat. No. 158388) according to the manufacturer’s instructions. Sequencing libraries were prepared using the TruSeq Nano DNA Sample Preparation Kit (Illumina), and paired-end sequencing (150 bp) was performed on an Illumina NovaSeq 6000 platform by Theragen Bio Inc. (Seoul, Republic of Korea).

### Variant calling and genotyping

Raw paired-end FASTQ files for each sample were aligned to the reference genome using BWA-MEM (v0.7.18) (Li, 2013) with default parameters. The resulting BAM files were sorted and indexed using SAMtools (v1.13) (Li et al., 2009). PCR duplicates were identified and removed using the MarkDuplicates tool in Picard (v.3.3.0) (REMOVE_DUPLICATES=true) (Broad Institute. Picard toolkit. http://broadinstitute.github.io/picard/). Subsequently, reads with a mapping quality score of at least 30 (Q ≥ 30) were retained using SAMtools (v1.13) for downstream analyses.

Variant calling for each sample was performed using GATK v4.6.1.0 HaplotypeCaller in GVCF mode (–ERC GVCF) (McKenna et al., 2010; Van Der Auwera et al., 2013). Individual GVCF files were merged using the CombineGVCFs module, followed by joint genotyping across all samples with GenotypeGVCFs to generate a multi-sample VCF file. As part of this process, bacterial contamination was assessed using Kraken2 (v2.1.3) (Wood et al., 2019)with a custom-built bacterial database constructed from the NCBI RefSeq bacterial genomes. The database was built using the kraken2-build utility with default parameters and cleaned prior to classification. One sample (JU993) exhibiting high bacterial contamination (≥40%) was excluded from all subsequent analyses (Supplementary Table S5).

Variant quality filtering was performed using the GATK VariantFiltration tool. In this study, filtering criteria were applied based on QUAL, QD, SOR, FS, MQ, MQRankSum, ReadPosRankSum, ExcessHet, and sequencing depth (DP). For DP, threshold values were defined to exclude extreme outliers based on the median depth distribution (median DP = 1,625), removing variants with depth less than one-fifth of the median or greater than five times the median. The following filtering expression was applied: (“QUAL < 30.0 || DP < 325.0 || DP > 8125.0 || QD < 2.0 || SOR > 3.0 || FS > 60.0 || MQ <= 40.0 || MQRankSum < -12.5 || ReadPosRankSum < -8.0 || ExcessHet > 10.0”). Variants that were not biallelic single-nucleotide polymorphisms (SNPs) were subsequently excluded, resulting in a final set of 3,204,155 high-quality biallelic SNPs.

Genotypes were assigned for all variants across all samples used for variant calling (n = 31). For each site, the most likely genotype was retained if the normalized genotype likelihood was ≥ 0.9; otherwise, the genotype was treated as missing. Finally, variant-level missingness and minor allele counts (MAC) were calculated, and variants with MAC ≤ 2 were removed. Only SNPs that passed all filtering steps were retained to generate the final SNP and genotype datasets for downstream analyses.

### Inference of Population Structure and Geographic Association

To infer the phylogenetic relationships among *Oscheius tipulae* strains, high-quality biallelic single-nucleotide polymorphisms (SNPs) were converted into a multiple sequence alignment using vcf2phylip (https://github.com/edgardomortiz/vcf2phylip) (Ortiz, 2019). The filtered multi-sample VCF file was transformed into a PHYLIP-formatted alignment, retaining only sites with a minimum allele count of four to reduce the impact of rare variants and missing data. Phylogenetic reconstruction was conducted under the maximum-likelihood framework using IQ-TREE version 3.0.1 (Minh et al., 2020), applying the GTR+Γ substitution model to account for heterogeneous substitution rates across sites. Node support was evaluated using 1,000 ultrafast bootstrap replicates. The inferred phylogeny was visualized in R using the ggtree (Yu et al., 2017) and treeio (Wang et al., 2020) packages.

Principal component analysis (PCA) was performed using smartpca v18140 from the EIGENSOFT package (Patterson et al., 2006), in which eigenvectors were computed from 31 diploid *O. tipulae* individuals. Ancestry inference was performed using ADMIXTURE v1.3.0 (Alexander et al., 2009)in an unsupervised framework, with the number of ancestral populations (K) ranging from 2 to 6. For each K, multiple independent runs were conducted, and the optimal K was determined based on the cross-validation error. To account for linkage disequilibrium (LD) and evaluate the robustness of the inferred population structure, SNP pruning was performed using PLINK v1.9 (Chang et al., 2015) with the --indep-pairwise 50 10 r² command, applying two LD thresholds (r² = 0.1 and r² = 0.8). PCA and ADMIXTURE analyses were then independently conducted on the unpruned dataset as well as on each LD-pruned dataset.

To assess the association between genetic similarity and spatial factors among *O. tipulae* isolates, Mantel tests were performed using pairwise genetic distances derived from genome-wide SNPs and two types of geographic distances (latitudinal differences and geographic distances). SNPs were LD-pruned as described above (r² = 0.8). Pairwise genetic similarity between all samples was then calculated using the identity-by-state (IBS) metric implemented in PLINK v1.9, generating a square similarity matrix. The IBS similarity matrix was converted into a genetic distance matrix by subtracting similarity values from 1. Latitudinal distances were calculated as the absolute differences in sampling latitude between all sample pairs, whereas geographic distances were computed based on the geographic coordinates of the samples. Each geographic distance matrix (latitudinal and geographic) was independently compared with the genetic distance matrix using Mantel tests. Correlations were evaluated using the vegan R package (Oksanen et al., 2001), employing Pearson’s correlation coefficient and 9,999 permutations to assess statistical significance.

### Population Relationships and Admixture Analyses

To further investigate population relationships and potential admixture among the three latitudinal groups, we performed admixture f3, outgroup f3, f4-statistic, and TreeMix analyses using genome-wide SNP data converted to EIGENSTRAT format. All analyses were conducted using the same filtered SNP dataset employed in the population genomic analyses described above.

Admixture f3 statistics were calculated using qp3Pop from the ADMIXTOOLS v7.0.1 package (Patterson et al., 2012). Tests were performed in the form f3(Target; Source1, Source2) to evaluate whether any of the three major lineages showed evidence of admixture between the other two lineages. Negative f3 values accompanied by significant Z-scores were interpreted as evidence of admixture.

Outgroup f3 statistics were also calculated using qp3Pop in the form f3(Outgroup; Population1, Population2) to quantify the amount of shared genetic drift between lineage pairs following divergence from the outgroup. Higher outgroup f3 values indicate greater shared evolutionary history between the two populations being compared.

Relationships among lineages were further evaluated using f4 statistics implemented in qpDstat (ADMIXTOOLS). Tests were performed using the outgroup lineage as the reference population to assess whether allele-frequency correlations were consistent with a strictly bifurcating tree or indicative of excess allele sharing among particular lineages. Statistical significance was assessed using block-jackknife standard errors and corresponding Z-scores.

To infer population relationships and potential migration events, we conducted TreeMix analyses using TreeMix v1.13 (Pickrell and Pritchard, 2012) with allele count data generated from the EIGENSTRAT genotype file. For each SNP, reference and alternative allele counts were calculated separately for the outgroup and each of the three latitudinal lineages. SNPs with missing allele counts in any population were excluded. TreeMix analyses were performed using a block size of 500 SNPs (-k 500), with the outgroup specified as the root of the tree. Models containing zero, one, and two migration edges (m = 0–2) were evaluated to examine whether migration events improved the fit of the inferred population graph.

### Genetic Diversity and Population Differentiation

Genome-wide nucleotide diversity (π), genetic differentiation (F_ST_), and sequence divergence (D_XY_) were estimated using Pixy (version 2.1.0) (Korunes & Samuk, 2021), which calculates unbiased population genetic statistics from all-sites VCF files while accounting for invariant sites. Analyses were performed using an all-sites VCF generated from the filtered genomic dataset.

Genome-wide summary statistics were subsequently obtained by aggregating 5-kb window-based estimates. Genome-wide π and D_XY_ were calculated as the total number of pairwise differences divided by the total number of pairwise comparisons across all windows, whereas genome-wide Hudson’s F_ST_ was calculated as the ratio of the summed numerator components to the summed denominator components across all windows. These calculations followed the approach recommended in the Pixy documentation.

### Local Adaptation and Selective Differentiation

To identify genomic regions showing excess population-specific differentiation beyond that expected from genome-wide background divergence, we quantified Population Branch Excess (PBE) following the median-scaling framework described by Yassin et al. (2016). Genome-wide differentiation was summarized in non-overlapping 5-kb windows. Within each window, mean F_ST_ values were calculated separately for each population pair. Pairwise genetic differentiation between populations was estimated using Hudson’s F_ST_ (Hudson et al., 1992). Windows containing fewer than 50 valid SNPs were excluded from further analysis. For each window, pairwise F_ST_ values were transformed into branch lengths using the standard population branch statistic (PBS) formulation (Yi et al., 2010):

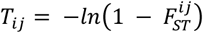

where *i*, *j*∈{SML,NML,LL}. Population-specific PBS values were then computed as:

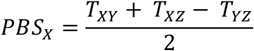

To control for genome-wide variation in background divergence, expected PBS values were estimated using a median-scaling approach. For each focal population X, the expected PBS was calculated as

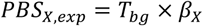

where T_bg_ is the background branch length (e.g., T_YZ_ for population X), and β_X_ is a scaling factor defined as:

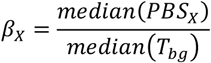

The Population Branch Excess for each window was then computed as:

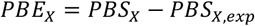

All windows were sorted by genomic coordinates, and PBE values were reported separately for each focal population (PBE_SML_, PBE_NML_, and PBE_LL_). Windows with high positive PBE values were considered candidate regions potentially shaped by population-specific positive selection and local adaptation.

To assess associations between environmental variables and genomic variation, we applied a Latent Factor Mixed Model (LFMM) (Caye et al., 2019; Frichot et al., 2013). LFMM accounts for unobserved confounding factors, such as population structure, while testing for associations between environmental variables and single nucleotide polymorphisms (SNPs).

Environmental variables were obtained based on the sampling locations (latitude and longitude) of each individual, using bioclimatic variables (bio1–bio19) and elevation extracted from the WorldClim database (Fick & Hijmans, 2017).

To correct for population structure, the number of latent factors (K) was estimated using the LEA R package, and the optimal K was determined based on cross-entropy (K = 4). LFMM analysis was performed using the lfmm2 algorithm, with each environmental variable analyzed independently. Associations between SNPs and environmental variables were tested using genomic control. P-values were adjusted for multiple testing using the Benjamini–Hochberg method (Benjamini & Hochberg, 1995), and SNPs with q-values < 0.05 were considered as environment-associated candidate SNPs.

To evaluate whether environment-associated candidate SNPs reflect signatures of selection, we examined their overlap with genomic regions corresponding to the top 1% of Population Branch Excess (PBE) values in each population.

For each PBE window, we calculated:

i. the total number of SNPs within the window, and
ii. the number of candidate SNPs identified by LFMM.

Using the total number of SNPs (N), the total number of candidate SNPs (K), the number of SNPs within each window (nᵢ), and the number of candidate SNPs within each window (kᵢ), we performed a hypergeometric test to assess whether candidate SNPs were significantly enriched beyond random expectation.

For each window, the expected number of candidate SNPs (expected = nᵢ × K / N) and fold enrichment were calculated. To account for multiple testing across windows within each environmental variable, p-values were adjusted using the Benjamini–Hochberg method. Windows with a false discovery rate (FDR) < 0.05 were considered as significantly enriched candidate regions.

Functional annotation was conducted for genes located within the significant regions. Gene sequences from the study species were mapped to orthologous genes in *C. elegans* using WormBase ParaSite BioMart (Howe et al., 2017).

Gene Ontology (GO) terms and functional annotations (descriptions) of the orthologous genes were then examined. By integrating environmental associations (LFMM), signatures of selection (PBE), and functional information, we identified candidate genes for local adaptation potentially driven by environmental factors.

## Data and resource availability

The sequencing data generated in this study have been deposited in the NCBI SRA under BioProject accession number PRJNA1464428. Previously published sequencing data analyzed in this study are available under NCBI SRA BioProject accession numbers PRJNA882448 (Dockendorff et al., 2022) and PRJNA974388 (Tintori et al., 2024).

## Acknowledgements

We thank Jiseon Lim and Jun Kim for collecting *O. tipulae* strains from Ulleung Island and Gwanak Mountain, Republic of Korea. We also thank Yeonpyeong High School and the Korea Foundation for Advanced Studies for assistance with nematode collection on Yeonpyeong Island. We are grateful to Professor Daehan Lee for helpful discussions on the manuscript.

## Author Contributions

D.H., D.S.L., and J.L. conceptualization; D.S.L. data curation; D.H. formal analysis; D.H. investigation; D.S.L. and J.L. supervision; D.H. visualization; D.H. writing—original draft; D.H., D.S.L., and J.L. writing—review and editing.

